# GABAergic neurons can facilitate the propagation of cortical spreading depolarization: experiments in mouse neocortical slices and a novel neural field computational model

**DOI:** 10.1101/2024.10.24.620012

**Authors:** Emre Baspinar, Martina Simonti, Hadi Srour, Mathieu Desroches, Daniele Avitabile, Massimo Mantegazza

## Abstract

Cortical spreading depolarization (CSD) is a wave of depolarization with local onset and extended propagation implicated in several pathological conditions. Its mechanisms have been extensively investigated, including our recent studies showing with experimental and computational approaches that the hyperactivity of GABAergic neurons’ can initiate migraine-related CSD because of spiking-generated extracellular potassium (K^+^) build-up. However, less is known about the role played by GABAergic neurons in CSD propagation. Here we studied mechanisms of CSD propagation, focusing on the role of GABAergic neurons, with experiments performed in mouse brain slices and with a new spatially extended neural field computational model. Experimentally, we induced CSD by applying brief puffs of potassium chloride (KCl) in somatosensory cortex slices from wild type and VGAT-ChR2-tdtomato mice, which specifically express the excitatory opsin channelrhodopsin (ChR2) in GABAergic neurons. We evaluated the role of GABAergic neurons in CSD propagation by modulating their activity with optogenetic illumination and their synaptic connections with pharmacological tools. We have developed the computational model to obtain realistic simulations of both initiation and propagation of CSD. It includes large populations of interconnected excitatory and inhibitory neurons, as well as the effect of extracellular ion concentrations on their features. We found that the decrease of the synaptic activity of GABAergic neurons can enhance CSD propagation, because of the reduction of the inhibitory synaptic weight, whereas their spiking activity can enhance CSD propagation because of extracellular K^+^ upload. However, differently than for CSD initiation, the latter effect is normally hidden by the action of GABAergic synaptic transmission. A reduction of GABAergic synaptic transmission, which can be observed in pathological states, can reveal the potentiating effect of the K^+^ upload induced by GABAergic activation. The neural field model that we implemented can generate accurate simulations of CSD, providing testable hypotheses on mechanisms, and can also be used for modeling other (patho)-physiological activities of neuronal networks.

## Introduction

Cortical spreading depolarization (CSD) is a wave of cellular depolarization that initiates locally and then slowly spreads across the cortex. It is characterized by an initial hyperexcitability with high frequency neural firing followed by prolonged neural silence which is induced by depolarization block of the firing [16, 31]. CSD is implicated in several pathologies: it is generated in normoxic conditions in migraine and epilepsy, whereas in stroke, traumatic brain injury and subarachnoid hemorrhage, it is generated in anoxic or hypoxic tissue. [16, 18, 29, 31].

CSD causes migraine aura and could induce migraine headache by sensitization of meningeal nociceptors [10, 16, 18, 28, 31]. Familial hemiplegic migraine (FHM), a genetic form of migraine with aura characterized by hemiplegia during the attacks, has become a model disease for investigating CSD mechanisms in migraine. In particular, FHM type 3 (FHM3) is caused by mutations of the

Na_V_1.1 voltage-gated sodium channel, encoded by the *SCN1A* gene [15, 29, 34]. Na_V_1.1 plays a key role in the excitability of GABAergic inhibitory neurons [39]. FHM3 mutations induce Na_V_1.1 gain of function, causing GABAergic neurons to fire at higher rate [4, 7, 8, 14, 29], which can lead to network hyperexcitability and CSD initiation. Therefore, GABAergic neurons can have a key role for triggering CSD initiation [11, 13, 25, 26]. However, although there is an extensive literature on the involvement of glutamate and glutamatergic neurons in CSD [31], much less is known about the influence of GABAergic neurons. In previous studies, we have investigated with experimental and modeling approaches the mechanisms linking hyperactivation of GABAergic neurons to CSD initiation, showing that the extracellular potassium build-up generated by the increased firing of GABAergic interneurons can lead to CSD [11, 13, 25, 26]. In particular, for investigating CSD initiation mechanisms, we have developed biophysical models of a pair of interconnected excitatory and inhibitory neurons, which included dynamics of ion concentrations [11, 13, 25]. However, the role of GABAergic neurons in CSD propagation has not been investigated yet. Here, we have performed experiments in mouse brain slices and developed a novel computational model to unveil it. In the experiments, we induced CSD by applying brief puffs of KCl in somatosensory cortex slices obtained from WT mice or VGAT-ChR2-tdtomato mice, which specifically express the excitatory opsin channelrhodopsin (ChR2) in GABAergic neurons [11]. We tested the role of GABAergic neurons in CSD propagation by modulating their activity with optogenetic illumination and their synaptic transmission with pharmacological tools.

Our computational model is a spatially extended neural field that can simulate the activities of large populations of glutamatergic (excitatory) and GABAergic (inhibitory) neurons. A part of previously proposed models for CSD dynamics considered neural population activity in terms of networks of single cells [12, 19, 21, 32], possibly by including metabolism dynamics [5, 22]. These models are high dimensional, and also computationally expensive due to the biophysical details in their cell descriptions. This limits the networks to a small number of cells compared to those that form biological networks. Global dynamics have been observed at the level of a cortical region, at the level of hundreds of population pairs where each population is composed of at least thousands of neurons. Therefore, these simulated networks are restricted to only local dynamics of CSD propagation. In some other models, CSD propagation was investigated on a larger scale but in terms of electrodiffusion, osmotic and ion variables, not in terms of variables describing the neural activity [17, 20, 24, 30, 35, 36, 37, 40]. This obscures the coupling between the neural activity and ionic changes in the extracellular matrix. Finally, CSD propagation dynamics were also modeled in terms of large number of populations in [23], however without making any distinction between excitatory and inhibitory neurons.

Neural fields model average population dynamics in a coarse-grained continuum limit. They are low dimensional compared to network models and with less biophysical details. Yet they can approximate closely the population dynamics. This allows to go far beyond the scale of local networks, scaling up to populations of thousands of neurons with synaptic interactions, and to generate global dynamics of CSD propagation. This motivates our choice for modeling approach. Moreover, our model considers neural interactions which are sensitive to changes in the extracellular potassium concentration, allowing to simulate the modulatory effects of potassium on the neural dynamics, which was not considered in classical neural field approaches [2, 38]. We calibrated and validated our model with the data obtained from the experiments, which in turn was used to test different hypotheses by performing specific simulations, also by simulating conditions that cannot be achieved experimentally.

Our modeling and experimental results show that the decrease of the synaptic activity of GABAergic neurons can enhance CSD propagation, because of the reduction of the inhibitory synaptic weight, whereas their spiking activity can enhance CSD propagation because of increased extracellular K^+^. The latter feature is revealed when GABAergic synaptic activity is reduced. Our neural field model proved to be important for generating testable hypotheses and better understanding mechanisms that modulate CSD propagation. Notably, its use can be extended to a variety of different other features of CSD and properties of interconnected neural populations.

## Results

### Induction of CSD and quantification of propagation speed in neocortical slices: comparison between WT and VGAT-ChR2-tdtomato mice

Since little is known about the role of GABAergic neurons in the propagation of CSD, we focused on this specific aspect in the experimental part of our study. To optogenetically control the activity of GABAergic neurons, we used VGAT-ChR2-tdtomato mice that specifically express channelrhodopsins (ChR2) in GABAergic neurons (see Materials and methods) and that we have already used in other studies [11]. We induced CSD in somatosensory cortical slices by applying brief puffs of 130mM KCl (Fig. 1) and we quantified CSD speed by measuring the spatial difference of the wave front propagating in layer II/III in images taken at different times and dividing the distance traveled by the elapsed time between the acquisition of the images (for each slice, we considered the average of 3 measurements: 4 images). We compared the properties of CSD propagation in slices obtained from VGAT-ChR2-tdtomato mice and WT littermates in control (without optogenetic illumination). CSD propagation speed was not different between the two groups (WT slices 2.13 ± 0.06 mm/min; VGAT-ChR2-tdtomato slices 2.35 ± 0.12 mm/min; Mann-Whitney test: *p* = 0.12) (Fig. 1F). Therefore, we pooled these data that we considered the control for the other experimental conditions used (Fig. 2). The local field potential (LFP) recorded with two extracellular electrodes placed at different positions showed the typical CSD DC shift when the wave-front crossed the zone in which the electrode was placed. We measured the CSD propagation speed also considering the distance between the two electrodes and the time lag of the onset of the DC shift. We found results that were similar to those obtained from the analysis of the IOS: WT slices 2.24 ± 0.08 mm/min; VGAT-ChR2-tdtomato slices 2.31 ± 0.11 mm/min: Mann-Whitney test: *p* = 0.14. For the subsequent analyses, we quantified the propagation speed considering the IOS; LFP recordings confirmed that the passage of the wavefront induced the characteristic DC shift associated with cortical spreading depolarization (CSD).

**Fig 1.**
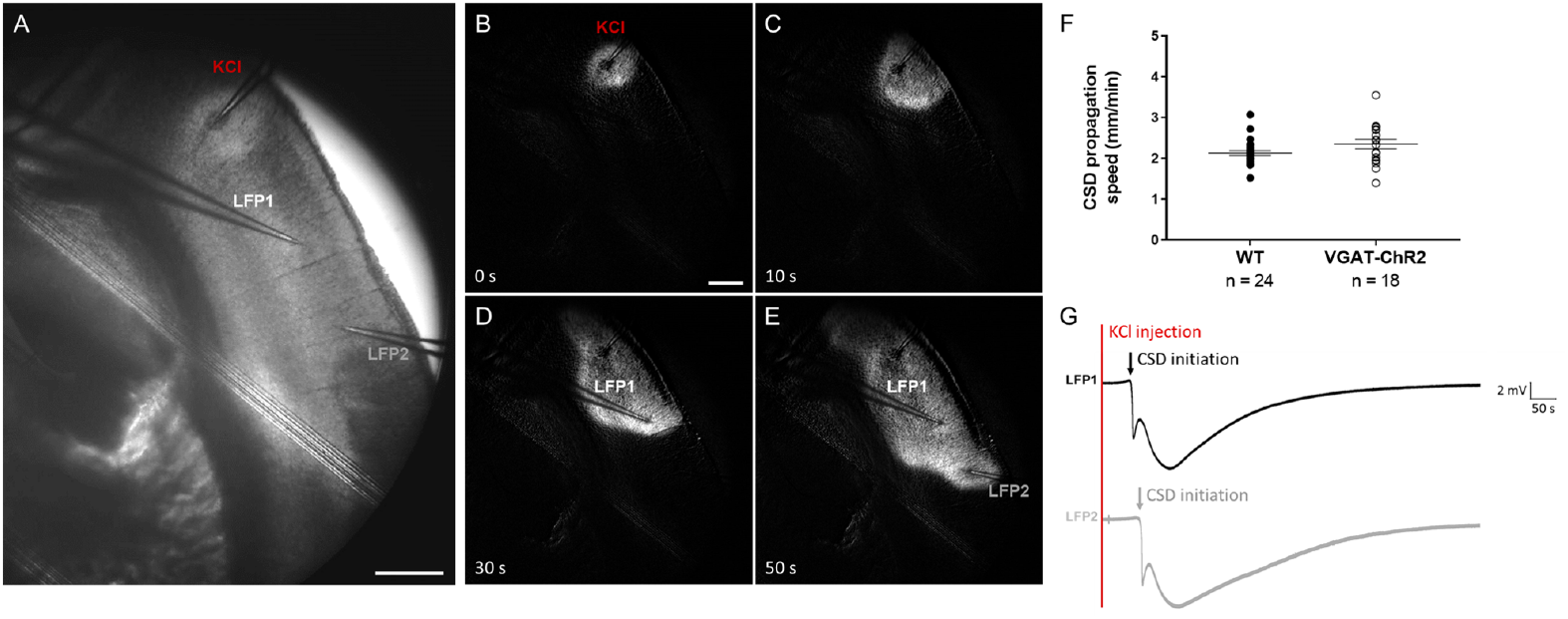
CSD experimentally triggered in somatosensory cortex mouse brain slices by puff application of 130 mM KCl. **A.** Raw near infrared transmitted light image of a representative coronal brain slice in which a pipette for applying 130 mM KCl and two extracellular electrodes for LFP recordings were placed in the somatosensory cortex. The image shows the initiation of CSD induced by a puff of KCl; Scale bar 0.5 mm. The other panels (**B-E**) show the same slice after image processing (subtraction of the background, optimization of the contrast) to highlight the intrinsic optical signal (IOS) during CSD initiation (KCl puff application, **B**) and propagation (**C-E**) at the indicated times; the white wave is the CSD that propagates in the cortex. Scale bar 0.5 mm. **F.** Comparison of the propagation speed measured in slices from WT mice and from VGAT cre − ChR2 lox mice (without optogenetic illumination), showing that there is no difference. **G.** LFP recorded during CSD propagation at the two locations indicated in **E** (LFP1 and LFP2), showing the typical CSD DC shift (note the delay of CDS initiation at LFP2 compared to LFP1).

**Fig 2.**
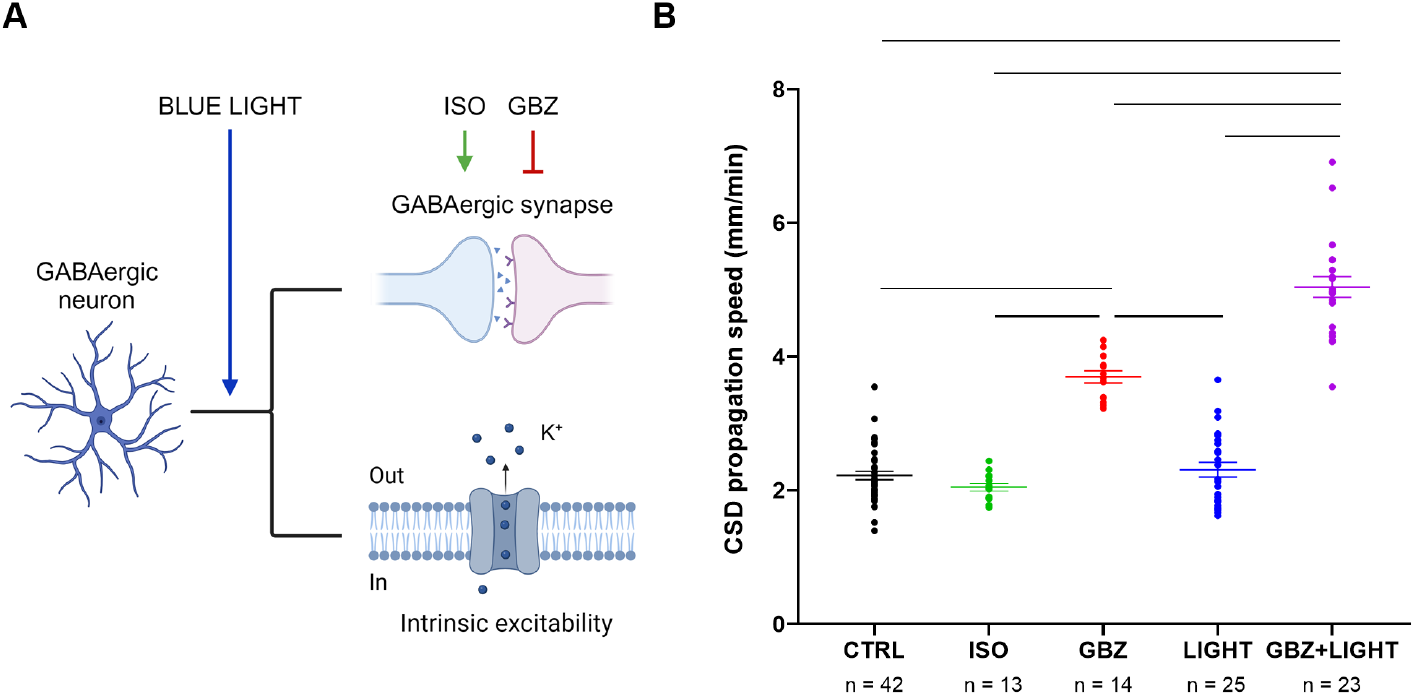
Effects of the modulation of GABAergic neurons activity and synaptic transmission on CSD propagation speed. **A:** Schematic representation of the effects of optogenetic illumination (BLUE LIGHT) on GABAergic neurons activity and synaptic transmission, and of the effects of isoguvacine (ISO) and gabazine (GBZ) on GABAergic synaptic transmission. Fig. 2A was created in BioRender. Mantegazza, M. (2025) https://BioRender.com/90to28g. B: CSD propagation speed in WT and VGAT-ChR2-tdtomato expressing slices (pooled) in control (CTRL, same data as in Fig. 1); in WT slices perfused with isoguvacine (ISO) or gabazine (GBZ); and in VGAT-ChR2-tdtomato expressing slices exposed to blue light in control (LIGHT) or with gabazine (GBZ + LIGHT). The number of slices (n) is indicated below the x axis. One-way ANOVA (*p <* 0.0001) with Tukey’s post hoc test; the bars indicate significant differences, all with *p <* 0.0001.

### GABAergic neurons have a dual role in CSD propagation: inhibition and enhancement

The activity of GABAergic neurons can lead to at least two actions with opposite roles: increased extracellular potassium because of the generation of action potentials, which can depolarize the network, and synaptic release of GABA with activation of GABA receptors, leading in general to an inhibitory action on the network. This inhibitory action in the latter is generated in particular by GABA_A_ receptors. We studied the effect on CSD speed of either activation or block of GABA_A_ receptors, as well as of optogenetic activation of GABAergic neurons, which leads to both an increase of action potential generation and of synaptic release of GABA with activation of GABA receptors (Fig. 2). We initially studied the effect of activation of GABA_A_ receptors: we triggered CSD with a KCl puff in brain slices perfused with the GABA_A_ receptor agonist isoguvacine (ISO), which did not modify CSD propagation (CTRL 2.22 ± 0.06 mm/min; ISO 2.05 ± 0.06 mm/min), as already reported [1]. In a separate set of experiments, we triggered CSD in slices perfused with the GABA_A_ receptor antagonist gabazine (GBZ), which induced a 1.7-fold increase in propagation speed (to 3.70 ± 0.09 mm/min). Then, we used optogenetic illumination with blue light (before or during CSD) to activate GABAergic neurons in slices expressing VGAT-ChR2-tdtomato. As we have already shown [11], this optogenetic stimulation selectively activates GABAergic neurons and, before the triggering of optogenetic-induced CSD, it does not lead to depolarization block. In the experiments included here, optogenetic-induced CSD was not observed, probably because they were performed during the latency period or CSD was triggered in a cortical area that was not imaged. Notably, we show here that the optogenetic activation of GABAergic neurons did not modify the propagation speed of KCl-induced CSD (LIGHT 2.31 ± 0.11 mm/min). Interestingly, the optogenetic illumination in VGAT-ChR2-tdtomato slices perfused with GBZ induced instead a 2.3-fold increase of propagation speed (GBZ + LIGHT 5.04 ± 0.15 mm/min), which was 1.4-fold larger than that induced by GBZ alone.

To confirm in a separate series of experiments the latter result, namely that the potentiating effect of optogenetic activation of GABAergic neurons on CSD propagation is revealed when GABA_A_ receptors are blocked, we illuminated with blue light VGAT-ChR2-tdtomato slices after inducing CSD with the KCl puff, in control conditions or in the presence of GBZ. Thus, we measured CSD propagation speed on the same slice before and during illumination with blue light, which was applied when the CSD wavefront was at around 700 *µ*m from the initiation site. In control conditions, blue light illumination did not modify propagation speed (2.20 ± 0.11 mm/min before blue light; 2.31 ± 0.11 mm/min during blue light; paired Student’s t-test: *p* = 0.35) (Fig. 3 A and B). In the presence of GBZ, blue light illumination induced instead a 1.3-fold increase in the propagation speed (4.07 ± 0.15 mm/min before blue light; 5.17 ± 0.15 mm/min during blue light) (Fig. 3 C and D). Therefore, we confirmed the results that we obtained comparing different slices, demonstrating that optogenetic activation of GABAergic neurons in conditions of intact GABAergic synaptic transmission is not sufficient to induce modifications in CSD propagation speed, but when coupled with GABA_A_ receptors blockade it produces a large increase. Taken together, these results indicate that the activation of GABAergic neurons has a double role of inhibition and enhancement on CSD propagation: the inhibitory role of GABAergic synaptic activity can mask the enhancement, in which the spiking activity and the increase in extracellular potassium should be key players.

**Fig 3.**
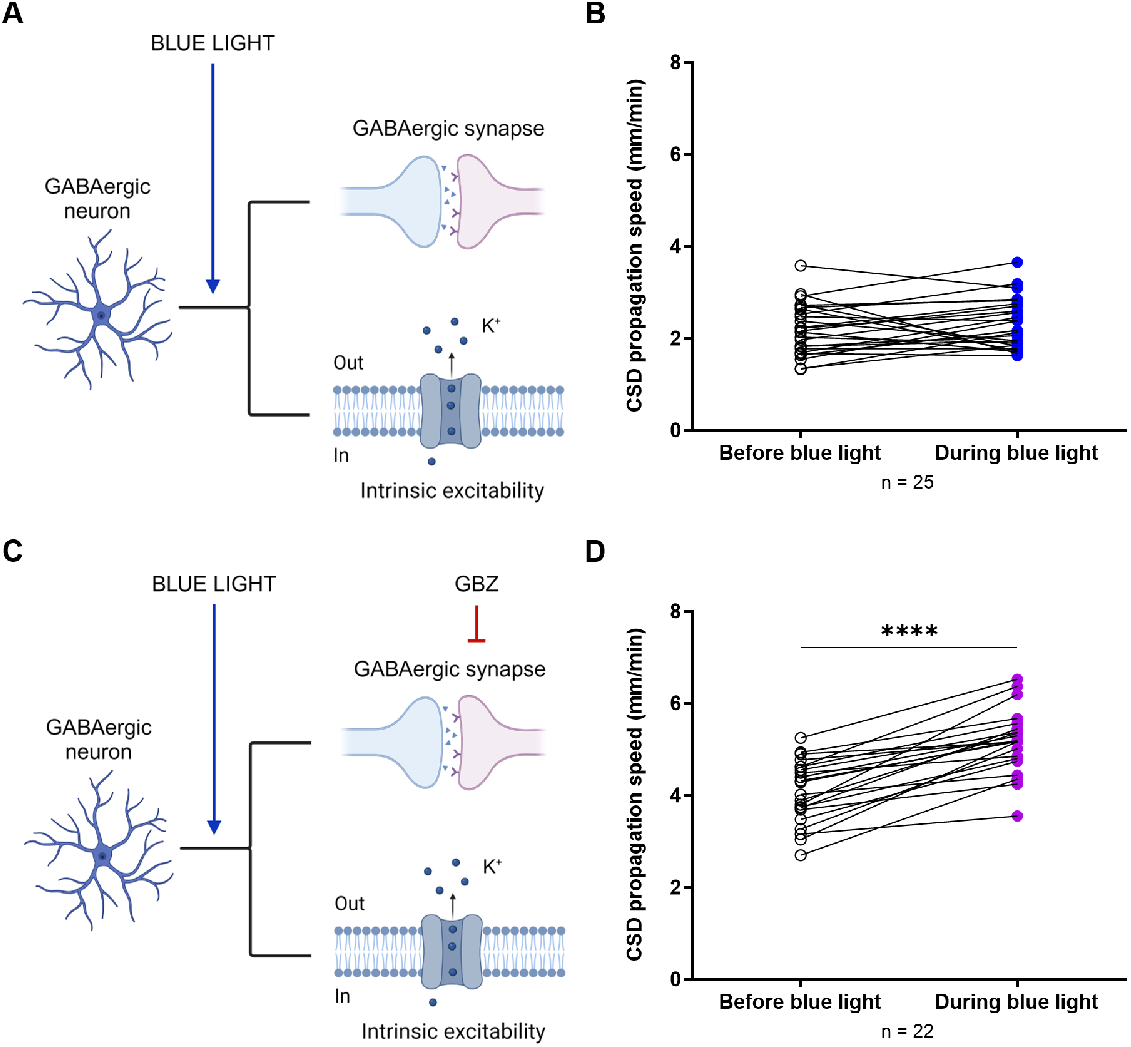
Effects of the optogenetic activation of GABAergic neurons on CSD propagation speed in the same slice. **A.** Schematic representation of optogenetic illumination effects on GABAergic neurons activity and synaptic transmission. **B.** CSD propagation speed in VGAT-ChR2-tdtomato slices before and during illumination with blue light. **C.** Schematic representation of the combined effects induced by optogenetic illumination and gabazine on GABAergic neurons activity and synaptic transmission. Fig. 3A and Fig. 3C were created in BioRender. Mantegazza, M. (2025) https://BioRender.com/90to28g. **D.** CSD propagation speed in VGAT-ChR2-tdtomato slices perfused with rACSF + gabazine before and during illumination with blue light. The number of slices (n) is indicated below the x-axis. Paired Student’s t-test: *p <* 0.0001.

### Neural field model

We have modeled CSD in terms of propagating waves described by average membrane potentials of excitatory (*v*_*e*_) and inhibitory (*v*_*i*_) neural populations, coupled to a reaction-diffusion equation which implements the dynamics of the extracellular potassium concentration (*k*). The equations of the model read:

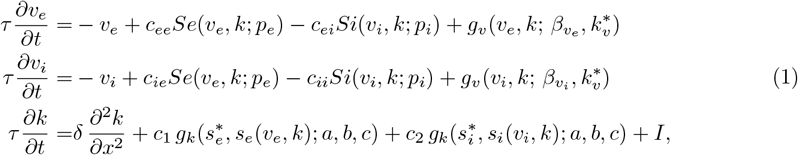

with the neural interaction terms

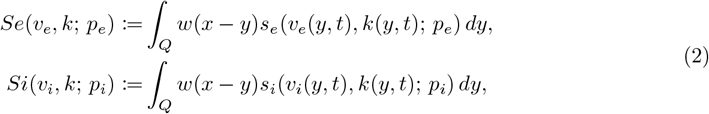

for excitatory and inhibitory interactions, respectively. Here *s*_*e*_ and *s*_*i*_ are the transfer functions of the excitatory and inhibitory populations, respectively (Eq. 3 in Material and Methods). They are convolved with 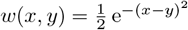 representing the connectivity kernel defined on a 1D propagation axis, which is represented by *Q* (Fig. 4). These kernels describe the neural interactions between the populations lying along the propagation axis (with weights *c*_*ee*_, *c*_*ie*_, *c*_*ei*_ and *c*_*ii*_ for connections between excitatory neurons, from excitatory to inhibitory, from inhibitory to excitatory neurons, and between inhibitory neurons, respectively). Here *g*_*v*_ is a generic function (Eq. 4 in Material and Methods) representing the effect of a humoral agent, in our simulations extracellular K^+^, on the neural activity expressed in terms of average membrane potentials: it implements any K^+^ related activity that affects the membrane potentials. Similarly, the effect of neural activity on the extracellular K^+^ concentration is introduced by the function *g*_*k*_ (Eq. 5 in Material and Methods), taking into account both excitatory and inhibitory neural activity. We denote by *I* the external input that models a focal CSD triggering stimulus, for example a localized potassium puff that is often used to induce CSD in laboratory experiments and that we have used in the experimental part of this study. The constant *δ >* 0 ensures the unit coherency between the diffusion and the drift in the potassium concentration equation. See Material and Methods for more details on the neural field model.

**Fig 4.**
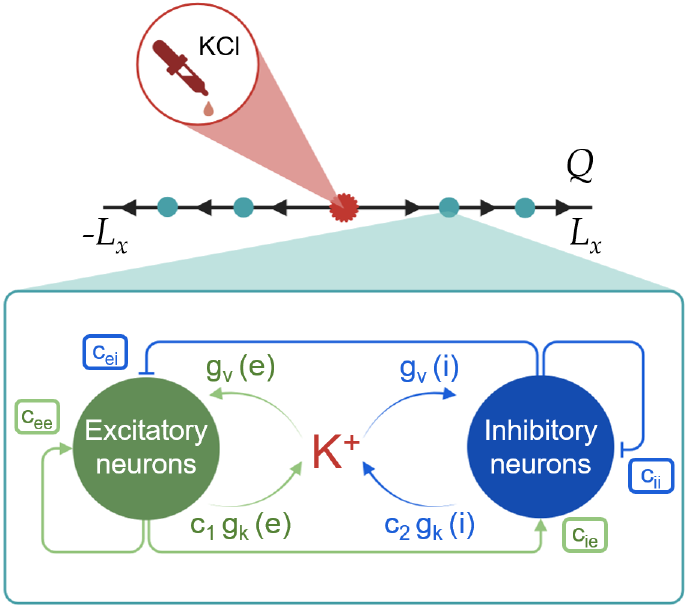
Illustration of CSD propagation over the propagation axis *Q* of the neural field model. The length of *Q* is 2*L*_*x*_. CSD initiation occurs at the source point marked in red. Each turquoise dot represents a microcircuit of interconnected excitatory and inhibitory neurons (schematically depicted in the diagram inset in green and blue, respectively). Here *c*_*ei*_ and *c*_*ee*_ model the connection weights of excitatory synapses on inhibitory neurons and of excitatory synapses on excitatory neurons (self-coupling), respectively. Similarly, *c*_*ie*_ and *c*_*ii*_ model the connection weights of inhibitory synapses on excitatory neurons and of inhibitory synapses on inhibitory neurons, respectively. The effect of excitatory and inhibitory neurons activity on the extracellular potassium concentration is represented by the functions *c*_1_ *g*_*k*_(*e*) and *c*_2_ *g*_*k*_(*i*), respectively. The influence of the extracellular potassium on excitatory and inhibitory neurons is described by the functions *g*_*v*_(*e*) and *g*_*v*_(*i*), respectively. Fig. 4 was created in BioRender. Mantegazza, M. (2025) https://BioRender.com/hfkou4f.

### Spatial extension

CSD is initiated locally in a relatively small area and then extensively spreads over the cortex. In our previous modeling and experimental frameworks, we extensively studied CSD initiation, focusing in particular on the role of the hyperactivation of GABAergic neurons [11, 13, 25]. However, those studies were not focused on the investigation of its propagation across the cortex, which is a main focus of the present study. The neural field in Eq. 1 constitutes one isolated unit of our model. This unit can simulate CSD initiation, but an interaction with other units is needed for simulating CSD propagation. Such an interaction requires a spatial extension of the isolated unit to a set of interacting neural fields.

For simplicity, we assume that CSD propagates horizontally and symmetrically. Although experimental neocortical CSD is not symmetric, because its propagation is faster in the superficial layers compared to the deep layers [41], the longitudinal propagation (within superficial or deep layers) is symmetric. In the experimental part of this study, we have quantified the speed of longitudinal CSD propagation in superficial layers. Consequently, our assumption allows us to consider a 1D propagation, along the propagation axis *Q*; see Fig. 4. Moreover, we assume that the neural field units lie along *Q* with equidistant spacing 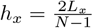, where the length of *Q* is 2*L*_*x*_ and *N* is the total number of units on *Q*.

We provide in Fig. 5A and 5B an example of CSD propagation over the spatial extension. Fig. 5A shows the space-time diagram of a CSD, defined as a wave of neural depolarization block that emerges from the population pair located at the center of the propagation axis *Q*. An external input, equivalent to a potassium puff, initiates the depolarization at the center, which then propagates as a wave towards the propagation axis boundaries horizontally and symmetrically with a constant velocity. On the front of the wave, the increasing potassium concentration gives rise to a depolarization and increased neural activity, then to a depolarization block, which propagates towards the periphery. Fig. 5B displays the plots of the propagating average membrane potentials of excitatory and inhibitory populations, as well as the increasing potassium concentration that expands towards the patch boundaries as the depolarization block propagates.

**Fig 5.**
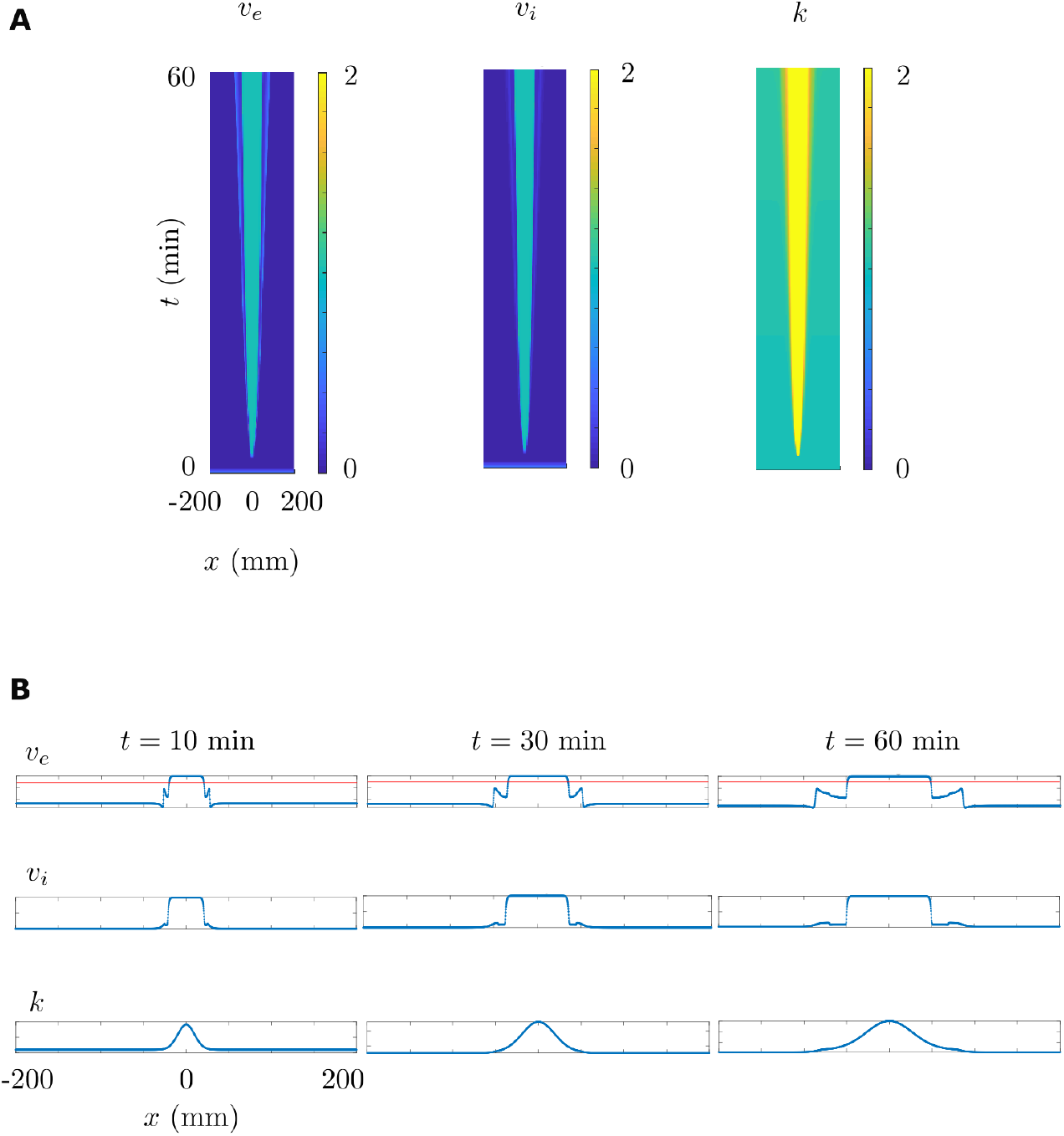
CSD propagation. **A.** Space-time diagrams of the propagation of excitatory and inhibitory population potentials, the expansion of potassium concentration in space and time. **B.** CSD propagation over the propagation axis *Q*. Top: Propagation in terms of average membrane potentials of the excitatory populations. Red horizontal line indicates the threshold detecting the wavefront of the propagating depolarization block. Middle: Propagation in terms of average membrane potentials of the inhibitory populations. Bottom: Expansion of the increasing potassium concentration towards the patch boundaries as the depolarization block propagates.

This neural field model can simulate both initiation and propagation of CSD and can be used to investigate specific properties of glutamatergic and GABAergic cortical populations, as well as their synaptic connections and the influence of extracellular potassium dynamics.

### Model-generated simulations of the experimental conditions give insights on mechanisms

We have used our computational model for generating simulations of the conditions that we have used in the experiments (Fig. 6). We considered ≈ 2.2 mm/min as the CSD speed in control conditions, as in the experiments that we performed.

**Fig 6.**
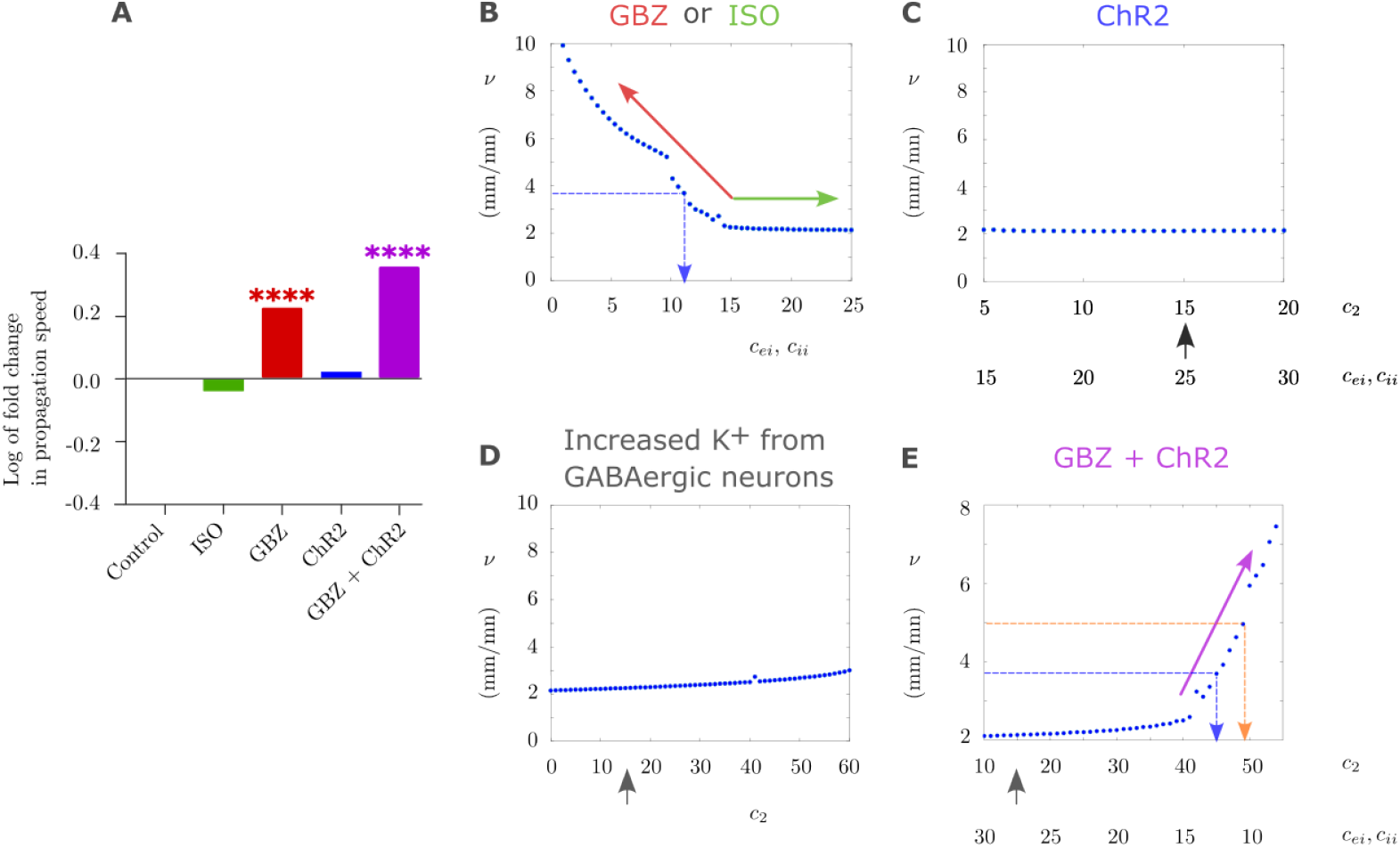
Simulation of the effect of the experimental conditions on CSD speed. **A.** Experimental results already displayed in Fig. 2, showed here as fold change. **B.** Effects of *c*_*ei*_, *c*_*ii*_ on CSD propagation speed. Increasing *c*_*ei*_, *c*_*ii*_ from the value that gives control propagation speed (mimicking ISO) results in a saturation in the speed, which corresponds to the plateau in the plot (green arrow). Decreasing *c*_*ei*_, *c*_*ii*_ (mimicking GBZ) results in an increase of the propagation speed (red arrow). The dashed blue line indicates the values of *c*_*ei*_, *c*_*ii*_ that produce a speed equal to that observed with GBZ in the experiments (3.7 mm/min). Here *c*_*ie*_ = *c*_*ee*_ = 1, *c*_1_ = *c*_2_ = 15 and *τ* = 0.15. **C.** Simulation of the modulatory effect of ChR2 activation with blue light on the propagation speed, resulting in an increase in the activity of the GABAergic population and, consequently, also of GABAergic synaptic transmission, which are modeled by increasing *c*_2_ and *c*_*ei*_, *c*_*ii*_, respectively, and that do not modify propagation speed. The black arrow indicates the value of *c*_2_ used in B. Here *c*_*ie*_ = *c*_*ee*_ = 1 and *τ* = 0.15. **D.** Effects of *c*_2_ (extracellular K^+^ generated from the GABAergic neurons) on the propagation speed. Increasing *c*_2_ results in a moderate increase in the propagation speed. Here *c*_*ei*_ = *c*_*ii*_ = 15, *c*_*ie*_ = *c*_*ee*_ = 1, *c*_1_ = 15 and *τ* = 0.15. The black arrow indicates the value of *c*_2_ used in B. **E.** Simulation of concomitant blocking of GABA_A_ receptors with GBZ and activation of ChR2, modeled by decreasing *c*_*ei*_, *c*_*ii*_ and increasing *c*_2_, respectively, in which the propagation speed shows a large increase (purple arrow). The dashed blue line indicates the value of the parameters that produce a speed equal to that observed with GBZ in the experiments (3.7 mm/min), the dashed orange line those that produce a speed equal to that observed with GBZ + blue light in the experiments (5 mm/min). The black arrow indicates the value of *c*_2_ used in **B**. Here *c*_*ie*_ = *c*_*ee*_ = 1 and *τ* = 0.15.

The first simulation was the effect of the activation of the GABA_A_ receptors during CSD, obtained experimentally by applying the GABA_A_ agonist ISO. We simulated this condition by increasing the parameters *c*_*ei*_, *c*_*ii*_, which model the connectivity weights of GABAergic synapses on excitatory neurons and of GABAergic synapses on inhibitory neurons, respectively (Fig. 6B). Increasing *c*_*ei*_, *c*_*ii*_ above the value that gives control speed (≈ 2.2 mm/min, obtained when *c*_*ei*_, *c*_*ii*_ = 15) does not increase CSD speed, which reaches a plateau (Fig. 6B). This is consistent with the results of the experiments and with a ceiling effect caused by the saturation of the GABAergic transmission during CSD. Then, we simulated the effect of the decrease of GABA_A_ receptors activation by decreasing *c*_*ei*_, *c*_*ii*_ (Fig. 6B), which we obtained in the experiments with the application of GBZ. In the experiments, blocking the GABA_A_ receptors results in an increase in the speed to 3.7 mm/min. In the simulations, a partial decrease in GABAergic synaptic weight (to *c*_*ei*_, *c*_*ii*_ = 11) is sufficient to reproduce this increase in the propagation speed (Fig. 6B). This seems counter-intuitive since a complete block of GABAergic receptors corresponds to *c*_*ei*_, *c*_*ii*_ = 0 and the increase in the speed for these vanishing synaptic weights is beyond the increase observed in the experiments. However, it may be due to the fact that this condition (*c*_*ee*_, *c*_*ii*_ = 0) completely blocks inhibition in the model, whereas in real neocortical circuits the block of GABA_A_ receptors, as obtained by the application of GBZ, does not completely block inhibition, because other sources of inhibition (other GABA receptors, neuromodulators, neuropeptides, etc.) can still be present.

In another simulation, we modeled the effect of ChR2 activation with blue light by increasing both *c*_2_ and *c*_*ei*_, *c*_*ii*_, which did not modify the propagation speed (Fig. 6C). This is similar to the results obtained from the experiments, in which the illumination with blue light did not modify the propagation. Mechanistically, the model shows that the effect of *c*_*ei*_, *c*_*ii*_ increase on propagation is in saturation for values that are larger than 15 (Fig. 6B), consistent with the lack of effect in Fig. 6C. A further parameter evaluated in Fig. 6C is *c*_2_, which simulates the effect of the K^+^ released by the activation of the GABAergic neurons. Fig. 6D shows that the increase of *c*_2_ has a moderate effect on propagation speed, which increases in particular when *c*_2_ values are higher than 20, consistently with the simulation displayed in Fig. 6C. In a further simulation, we modeled the effect of the concomitant block of GABA_A_ receptors with GBZ and activation of ChR2-expressing GABAergic neurons with blue light by decreasing *c*_*ei*_, *c*_*ii*_ and increasing *c*_2_, respectively (Fig. 6E). In particular, considering that the GABAergic synaptic weight corresponding in the simulations to the presence of GBZ is *c*_*ei*_, *c*_*ii*_ = 11 (Fig. 6B), we set in this simulation *c*_2_ so that with values of *c*_*ei*_, *c*_*ii*_ close to 11 (i.e., 10.5), the propagation speed was 5 mm/min, which is the value observed in the experiments when both GBZ and blue light were applied (see Fig. 2 and 3). We found that with this setting the corresponding value of *c*_2_ was 49 and the simulation reached to propagation speeds that were higher than those induced by GBZ alone (Fig. 6B). Interestingly, this *c*_2_ value is higher than that corresponded in the simulation of Fig. 6C to the speed observed experimentally. Thus, the model shows that higher extracellular K^+^ can be generated at the CSD wavefront in the condition in which there is both GBZ application and illumination with blue light, compared with the same blue light illumination without GBZ (Fig. 6C). This is consistent with a larger accumulation of extracellular K^+^ induced by the activation of the GABAergic neurons in this condition.

## Discussion

We performed experiments in brain slices and developed a novel neural field computational model to investigate features of CSD, which is a wave of neural hyperexcitability followed by prolonged depolarization of the network that initiates locally and then spreads across the cortex [16, 31]. Normoxic CSD is a proposed pathological mechanism of migraine. We used here our computational model principally for studying the role of GABAergic neurons in normoxic CSD propagation, integrating experimental results and generating testable hypotheses. CSD propagation is an important feature that we did not model in detail in our previous experimental and computational studies of CSD mechanisms, which were mainly focused on mechanisms of initiation [11, 13, 25, 26]. Importantly, differently from other models of CSD, our neural field model allows to simulate large populations of interconnected GABAergic and glutamatergic neurons.

The model is based on a Wilson-Cowan-Amari type framework [2, 38], in which an excitatory-inhibitory neuron population pair is coupled to extracellular K^+^ concentration. The extracellular K^+^ concentration is described in terms of a reaction-diffusion equation. The model has several novelties compared to a classical Wilson-Cowan-Amari model. First, in the firing rate transfer function, we include not only the firing activity, but also the extracellular K^+^ concentration, which is dynamic and depends on the firing activity. In addition, a transfer function from K^+^ to firing rate is introduced as an input to firing activity. We found in previous studies that this interaction between the firing activity and the extracellular K^+^ concentration is key for the CSD initiation induced by hyperactivation of GABAergic interneurons [11, 13, 25, 26]. Second, in contrast to the models that we previously developed [11, 13, 25, 26], our present model can simulate not only CSD initiation, but also CSD propagation. Third, parameters that influence the propagation speed can be easily adjusted for evaluating their effect in different conditions. For example, it provides control on the connection weights and contribution weights of different populations to extracellular K^+^ accumulation. Fourth, the time scale appearing in Eq. 1 can be tuned according to the experimental conditions, such that realistic propagation speeds can be obtained in the simulations. This provides a strong flexibility to our model, in the sense that it can be tuned to different experimental conditions, allowing to study mechanisms and generate hypotheses about CSD, but also to other (patho)-physiological activities of neuronal networks.

For future studies, our model can be extended to also include astrocytes, which are involved in numerous activities, including spatial K^+^ buffering and constitute an active research area in migraine and epilepsy [6, 33, 42]. They can be inserted in our model equations (1) as a third cell population coupled to the excitatory and inhibitory populations.

Regarding the biological mechanisms, in the present work, we focused on the role of hyperactivation of GABAergic neurons in CSD propagation, which could result from hemiplegic migraine mutations of the *SCN1A* gene, causing gain of function of Na_V_1.1 voltage-gated sodium channels leading to hyperexcitability of GABAergic neurons [11, 13, 25, 26]. In particular, we aimed at disclosing the relative effect of neural excitability and related increase in extracellular K^+^ vs synaptic transmission. We experimentally studied the role of GABAergic neurons in CSD propagation by increasing their intrinsic excitability with optogenetic stimulations (which increases both firing and synaptic transmission) and modulating the effect of their synaptic transmission with GBZ and ISO, which are specific inhibitor and activator of GABA_A_ receptors, respectively. We observed that a decrease of GABA_A_ receptor activation, and therefore disinhibition, facilitates CSD propagation by increasing its speed, whereas their pharmacological activation has no effect. These results confirm findings already reported [1, 11, 16, 31]. In particular, Aiba and Shuttleworth proposed a ceiling mechanism for the lack of effect on CSD propagation of the activation of GABA_A_ receptors, which could be already saturated by the GABA released at the wave front of CSD [1]. Importantly, the effect of GABAergic neurons’ excitability on CSD propagation has instead never been studied before. Here, we activated GABAergic neurons by applying optogenetic illumination either before or after the triggering of CSD by KCl pufs. In both cases, the illumination did not modify the properties of CSD propagation. However, a facilitatory effect leading to increased CSD speed was observed in disinhibited slices that were pre-treated with GBZ (Fig. 6A). These results show that increased

GABAergic neurons’ intrinsic excitability can not only induce CSD initiation, as we have previously shown [11, 13, 25, 26], but can have facilitatory effects also on the propagation in the condition of disinhibition, probably through an increase of extracellular K^+^. Differently than for initiation, this facilitatory effect is hidden by GABAergic synaptic transmission.

Our computational model generated detailed simulations of CSD and reproduced our experimental observations. We used the computational model to better identify the mechanisms that lead to the features of CSD propagation which we have observed experimentally, in particular to discern the contribution of GABAergic synaptic transmission and intrinsic excitability, which were modulated in the computational model by modifying connection weights and contribution weights of the populations to extracellular K^+^ accumulation, respectively. Notably, by providing control also over parameters that can be difficult to control in an experimental setting, the model can allow investigating the conditions and their related mechanisms that are difficult to study experimentally. In particular, we reproduced the ceiling effect in the activation of GABA_A_ receptors during CSD propagation, which was induced by the saturation of the receptors at the CSD wave front (Fig. 6B). Moreover, we have reproduced the potentiating effect observed in a disinhibited cortical network (Fig. 6C) as well as the lack of effect of the activation of GABAergic neurons on CSD propagation in physiological conditions (Fig. 6D). Importantly, the simulations suggest that the potentiating effect is caused by a larger release of K^+^ from GABAergic neurons at the CSD wave front when the network is disinhibited (Fig. 6E). This could be caused by a larger and faster-onset of hyperexcitability in the initial phase of CSD (before the depolarization), induced by the reduced inhibition in the network. Notably, disinhibition may be present in pathological conditions, thus this may be a relevant condition in some pathological states. Furthermore, the simulations suggest that in our experimental conditions the block of GABA_A_ receptors with GBZ at the CSD wave front is not complete, because CSD in the presence of GBZ should be faster if this was the case (Fig. 6B).

In conclusion, we have implemented a neural field computational model that can generate accurate simulations of CSD, providing testable hypotheses on mechanisms, and that can also be used for modeling other (patho)-physiological activities of neuronal networks. Our modeling and experimental results show that GABA_A_ receptors can modulate CSD propagation speed, but they are probably already engaged at saturation during CSD propagation. Activation of GABAergic neurons can promote CSD propagation by spiking induced extracellular K^+^ upload, but this effect is hidden by GABAergic synaptic transmission. Reduced GABAergic synaptic transmission in pathological conditions may disclose the potentiating effect of GABAergic neurons’ spiking.

## Materials and methods

### Ethics statement

Experiments were carried out according to the ARRIVE guidelines and the European directive 2010/63/UE, and approved by local (the “Comité Institutionnel d’éthique pour l’Animal de Laboratoire - CIEPAL” of the Côte d’Azur and the “Structure chargée du Bien Etre des Animaux - SBEA” of the Institute of Molecular and Cellular Pharmacology - IPMC) and national (Minist’ere de la Recherche, France) ethical committees (approval PEA 362/APAFIS 7848-2016112114249820v3).

### Animals

All mice were housed under standard conditions (22–24°C, 40– 50% humidity, 12h dark/light cycle) with access to food and water *ad libitum*. After weaning (post-natal days P21–23), mice were group-housed with same sex littermates of mixed genotype (n = 3–5*/cage*) in standard polycarbonate cages containing plastic houses, wood sticks and nesting material.

The mouse lines were obtained from Jackson Laboratory (JAX) in the C57BL/6J genetic background. As previously described [11], we generated double hemizygous transgenic mice VGAT-hChR2(H134R)/tdtomato (VGAT-ChR2), which specifically express ChR2-H134R/tdtomato in GABAergic neurons, by crossing hemizygous females loxP-STOP-loxP-hChR2(H134R)-tdTomato (Ai27D, B6.Cg-Gt(ROSA) 26Sortm27.1(CAG-COP4*H134R/tdTomato)Hze/J; JAX catalog 012567) [27] with transgenic males (to avoid off-target Cre expression in the female germline) hemizygous for the Viaat-Cre transgene (Cre recombinase expression driven by the vesicular GABA transporter, VGAT, promoter) (B6.FVB-Tg(Slc32a1-Cre)2.1Hzo/FrkJ; JAX catalog 017535), line 2.1 [9]. Experiments were performed with 4-6 weeks old mice from both sexes.

### Preparation of brain slices

Mice were anesthetized with isoflurane, the brain quickly removed and acute 400 − *µ*m thick coronal slices of the somatosensory cortex were obtained with a Leica VT1200 vibratome as in [11, 41]. Slices were then stored before the beginning of the recordings at 32°C for one hour in an incubation chamber filled with artificial cerebrospinal fluid (ACSF), which contained (in mM): 129 NaCl, 3 KCl, 1.6 CaCl_2_, 1.8 MgSO_4_, 1.25 NaH_2_PO_4_, 21 NaHCO_3_ and 10 glucose, saturated with 95% O_2_,5% CO_2_.

### Induction of CSD, intrinsic optical signal analysis and local field potential recordings

Single slices were transferred to the recording chamber and perfused with recording ACSF (rACSF) at 32°C, containing (in mM): 129 NaCl, 3.5 KCl, 1 CaCl_2_, 0.5 MgCl_2_, 1.25 NaH_2_PO_4_, 21 NaHCO_3_ and 25 glucose, saturated with 95% O_2_, 5% CO_2_, as in [11, 41].

Slices were visualized using infra-red differential interference contrast (DIC) microscopy with an Eclipse FN1 microscope (Nikon, Japan) equipped with a CCD camera (CoolSnap ES2, Photometrics, USA), filter cubes for visualization of fluorescent proteins (Semrock, USA) and a filter for optogenetic 475nm blue light illumination (FF02-475/50; Semrock, USA). Acquisitions of large fields of view were obtained with a 4X objective and a 0.35X camera adapter lens.

CSD was induced as previously described (Zerimech et al., 2020; Chever et al., 2021). Briefly, short puffs of KCl (130 mM) and Fastgreen (0.1% Sigma-Aldrich) were applied in the superficial cortical layers (L2-3) with a glass micropipette (2-4 MΩ), connected to an air pressure injector (PV820 Pneumatic Picopump, WPI, USA).

Extracellular DC local field potential (LFP) was recorded with a Multiclamp 700B amplifier (CV-7B headstage), a Digidata 1440A acquisition board and pClamp 10.3 software (Molecular Devices, USA). DC extracellular field potential recordings were performed using Ag/AgCl electrodes and two borosilicate glass micropipettes filled with rACSF, placed in Layer 2-3 about 1 (LFP1) and 1.5mm (LFP2) away from the site of CSD induction (Fig. 7).

**Fig 7.**
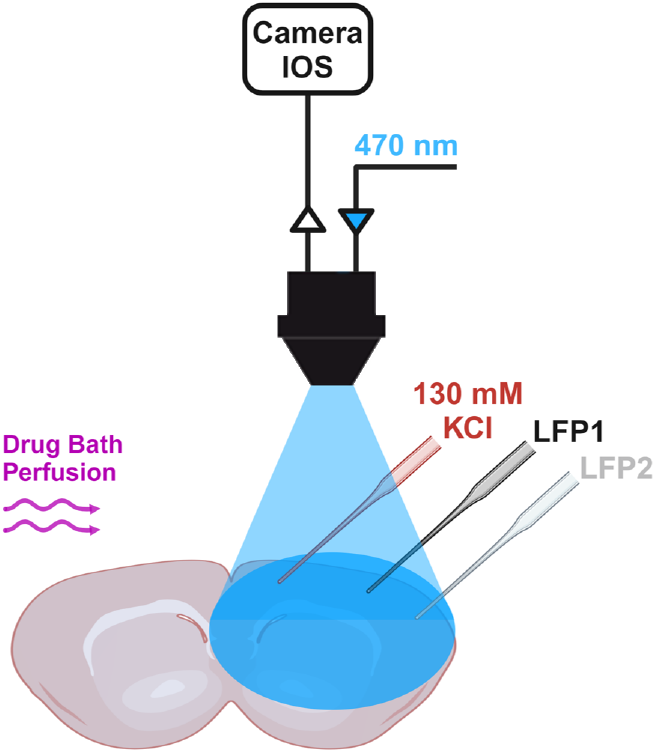
Experimental setting. CSD was induced in somatosensory cortex slices by applying a puff of KCl 130 mM. CSD waves were visualized by near infrared intrinsic optical imaging (IOS) with a CCD camera. Two micropipettes, placed about 1 mm (LFP1) and 1.5mm (LFP2) away from the site of CSD induction, were used to record LFPs. Drugs to modulate GABAergic neurons’ synaptic activity were applied to the bath. The activity of GABAergic neurons, which expressed ChR2, was induced by optogenetic illumination with blue light (470 nm). Fig. 7 was created in BioRender. Mantegazza, M. (2025) https://BioRender.com/6zlclb8.

CSD waves visualized by intrinsic optical imaging (IOS) were processed and analyzed with ImageJ-Fiji as previously described in [11, 41]. CSD propagation speed was quantified by measuring the spatial difference of the wave front propagating in layer 2-3 in images taken at different times and dividing the distance traveled by the elapsed time between the acquisition of the images (for each slice, we considered the average of 3 measurements: 4 images). LFP recordings were used to confirm that the passage of the wave-front induced the typical CSD DC-shift and to confirm the quantification of the propagation speed obtained with the IOS imaging.

### Pharmacology

Slices were perfused with rACSF + 15 *µ*M Gabazine or with rACSF + 10 *µ*M Isovugacine starting 10 minutes before inducing CSD. Drugs were purchased from Sigma-Aldrich.

### Optogenetics

Activation of ChR2 was obtained illuminating brain slices through the 4x objective (continuous illumination), as in [11]. A white light source (130 W mercury lamp, Intensilight, Nikon, Japan) was connected to the epi-illumination port of the microscope with a light guide containing a 420 nm UV blocker filter (series 2000, Lumatec, Germany); the white light was filtered with a 475/50 filter (Semrock) placed in the optical path and delivered to the objective with a FF685-Di02 dichroic beamsplitter (Semrock). The area of illumination was 38.5 mm^2^. The blue light power density measured with a power meter (Ophir Photonics, USA) was 2.8 mW/mm^2^ at the slice surface (photodetector placed at the level of the slice). According to the type of experiment, slices were illuminated either 3-4 s before CSD induction or after CSD induction, when the CSD wave-front was at around 700 *µ*m from the initiation site. This optogenetic stimulation selectively activates GABAergic neurons and, before the triggering of optogenetic-induced CSD, it does not lead to depolarization block, as shown in [11]. Here, we have not observed optogenetic-induced CSD, probably because the experiments were performed during the latency period or CSD was triggered in a cortical area that was not imaged.

### Statistics

Mice were used after genotyping and littermates were negative controls. Experimenters were not blinded to the genotype; animals were randomized (http://www.randomizer.org) within each experimental group. We used at least 3 animals per each experimental condition. The number of slices used (n) for each condition is indicated in the results. Data are presented as mean ± SEM.

Statistical tests were performed with GraphPad Prism 9. The Shapiro-Wilk test was used to assess normality and the Brown-Forsythe test was used to evaluate homogeneity of variances. For independent groups with normal distributions, we used unpaired student t-test. When variances were unequal, the Welch correction was applied. Paired t-test was used to compare two paired groups with normal distribution. Non-parametric Mann-Whitney U test was used when distributions were not normal. To compare three or more groups, we used One-way ANOVA, followed by Tukey’s post hoc test for multiple comparisons. Significance threshold was set at *p* = 0.05.

### Model

#### Firing rate transfer function

In a classical neural field, low inhibitory activity gives rise to high excitatory activity and high inhibitory activity gives rise to low excitatory activity. In our framework, the firing rate transfer function takes into account the generalized excitatory activity that distinguishes very high inhibitory activity from high inhibitory activity. It is defined for a population *α* ∈ {*e, i*} as

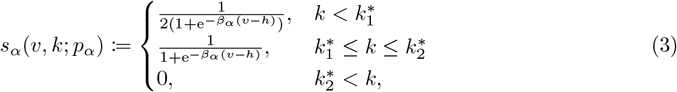

where 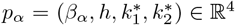 denote the parameters. Here *β*_*α*_ determines the sharpness of the nonlinearity of *s*_*α*_, and *h* expresses the value of the half response voltage. Finally, 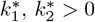 denote the potassium thresholds.

The corresponding output of the transfer function to a very high inhibitory activity is a low excitatory activity or a silent phase. This property of the transfer function is due to a threshold effect. Unlike the classical neural field setup, where the transfer function depends only on the neural activity, in our framework the transfer function also depends on the time-dependent potassium concentration *k*; see Fig. 8. Consequently, the same neural activity can produce different transfer function outputs depending on the potassium concentration in the extracellular matrix. In other words, we observe a threshold effect when the potassium concentration is very high, i.e., when ^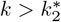^. The transfer function has the typical nonlinear sigmoidal behavior for ^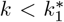^and ^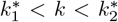^, where the maximum value of the transfer function is higher for the latter. The former corresponds to a low potassium concentration and thus a low inhibitory activity regime with 1*/*2 as the maximum firing rate. The latter corresponds to a high potassium concentration and thus to a high inhibitory activity with 1 as the maximum firing rate. A neural regime above the threshold with ^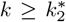^ corresponds to a very high inhibitory activity, in which the potassium concentration is too high. This results in a low excitatory firing rate for all *v*_*α*_ values and generates the depolarization block that marks the onset of CSD and triggers it; see Fig. 8.

**Fig 8.**
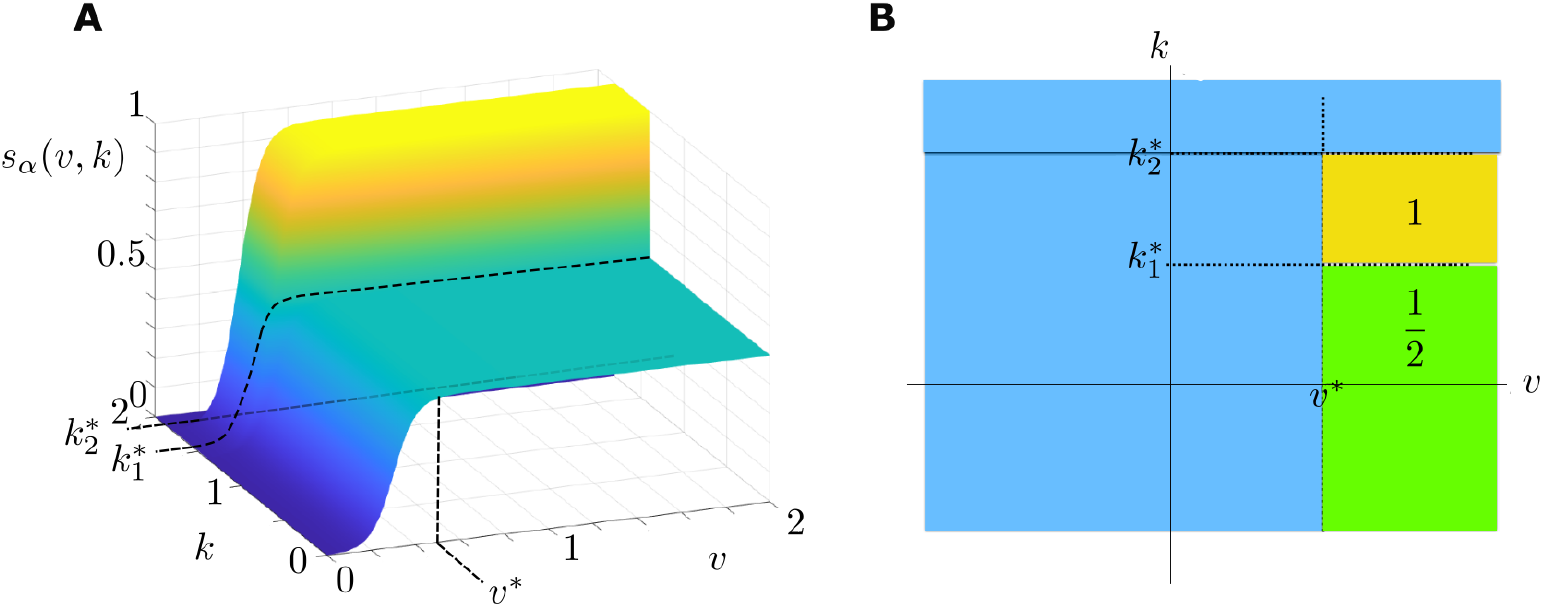
Transfer function plots. **A.** Transfer function *s*_*α*_ of a population *α* ∈*{e, i}* with respect to the firing activity *v* and potassium concentration *k*. **B.** Top view of the maximum and minimum values of the transfer function. The nonlinear regions are ignored.

This choice of the transfer function with three potassium levels is motivated by the neural activity appearing when the neural excitability is perturbed due to SCN1A mutation. This mutation is considered to be at the origin of CSD observed in FHM3 type migraine. As a result of this mutation, the neurons become highly excitable as the potassium level increases in the extracellular matrix. This is the phase of propagating hyperactivity, which marks the wavefront of CSD. This increase in potassium level eventually causes the neurons to become saturated. This is the phase of the spreading depolarization block observed in CSD. In this context, the three potassium levels in the activation function represent three regimes: (i) Normal neural activity regime 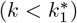 where the maximum value of sigmoid in *s*_*α*_ is 1/2. (ii) Hyperactivity regime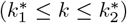, which appears as potassium level increases further in the extracellular matrix. This corresponds to a sigmoid but now with a maximum activity value 1, which is the double of that of the normal activity regime. (iii) Depolarization regime. Here, the neural activity stops. The neurons are saturated due to the very high potassium level. This is observed as the appearance of the depolarization block.

The code of the model implementation is available and accessible in [3].

#### Potassium to neural activity transfer function

We consider potassium-related effects on the population activity via *g*_*v*_, which is a nonlinear function given by

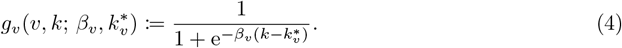

Here *β*_*v*_ *>* 0 and ^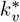^ determine the sharpness of *g*_*v*_ and *k* denotes the threshold. Note that *g*_*v*_ is a generic function which models all potassium-related population activity, with the exception of potassium build-up due to the potassium currents resulting from neural spikes. This latter type of activity is explicitly supplied to the system via the firing rate transfer function *s*_*α*_. See Fig. 9 for a simplified plot of *g*_*v*_.

**Fig 9.**
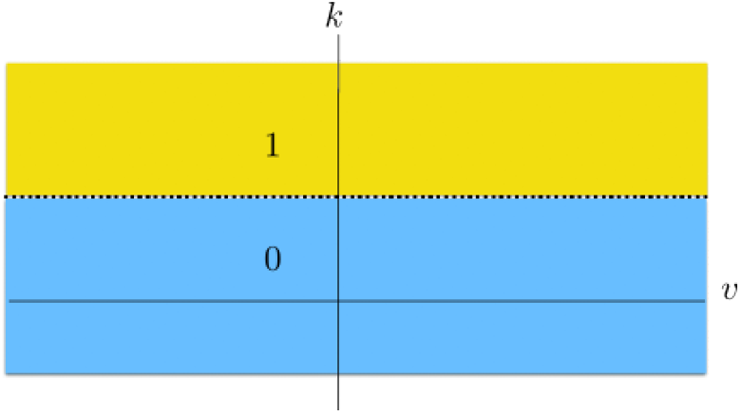
Potassium to rate transfer function *g*_*v*_ with respect to the population firing rate *v* and potassium concentration *k*. Regions of the maximum and minimum values are represented by the yellow and blue colors, respectively.

Parameter 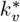 termines the activity regime inducing the CSD ignition and propagating depolarization block, see Fig. 10.

**Fig 10.**
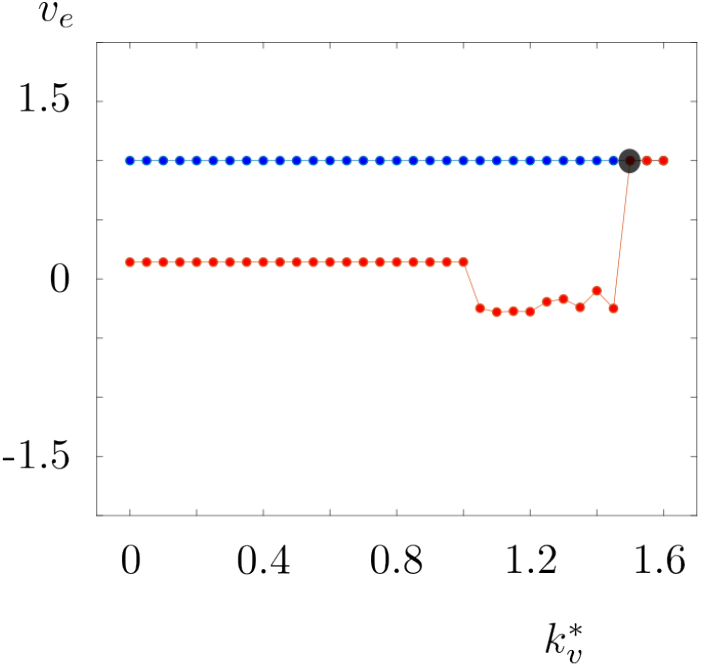
Bifurcation diagram of one node of the model with respect to ^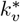^. Blue and red dots show the maximum and minimum values of *v*_*e*_ during the course of simulation. We see that as we decrease ^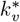^ to the values lower than 1.5, the system goes through a bifurcation (black dot) and switches to a regime with propagation waves. Here *c*_*ie*_ = *c*_*ee*_ = 1, *c*_*ei*_ = *c*_*ii*_ = 15, *c*_1_ = *c*_2_ = 15 and *τ* = 0.15.

#### Firing rate to potassium transfer function

The nonlinear function *g*_*k*_ models the effects of the interactions between the potassium concentration and the excitatory-inhibitory neural activity, on the potassium concentration. We define *g*_*k*_ as follows:

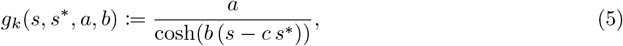

with parameters *a >* 0, *b, c* ≥ 0, *s* = *s*_*α*_(*v, k*) and 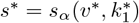 where *v*^*∗*^ > *h*.

#### CSD propagation velocity

The propagation velocity *ν* is obtained based on

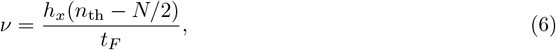

where *h*_*x*_ represents the physical distance between two adjacent indices lying on *Q*; see Fig. 4. Moreover, *n*_th_ and *t*_*F*_ denote the index of the outmost excitatory population having an average potential over the preset threshold shown by the red horizontal line in Fig. 5B and the final time, respectively.

## Code availability

The code of the model implementation is accessible in [3].

## Acknowledgments

We thank Emilie Bonnet (Institute of Molecular and Cellular Pharmacology, CNRS UMR 7275 Inserm U1323 and University Côte d’Azur) for her skillful technical support for the management of mouse colonies. This work was supported by the Investissements d’Avenir-Laboratory of Excellence *Ion Channels Science and Therapeutic* (grant LabEx ICST ANR-11-LABX-0015-01 to M. Mantegazza) and University Côte d’Azur (grant IDEX Jedi ANR-15-IDEX-01 to M. Mantegazza). M. Simonti was a PhD student of the *École Doctorale 85* of the University Côte d’Azur and received a LabEx ICST PhD fellowship. The laboratory of M. Mantegazza is a member of the *Fédération Hospitalo-Universitaire* FHU-INOVPAIN consortium and of the Interdisciplinary Institute for Modeling in Neuroscience and Cognition of the University Côte d’Azur (Neuromod Institute). M. Desroches was supported by the grant PID2023-146683OB-100 funded by MICIU/AEI/10.13039/501100011033 and by ERDF, EU. Additionally, M. Desroches acknowledges the Ikerbasque Foundation and the Basque Government through the BERC 2022-2025 program and by the Ministry of Science and Innovation: BCAM Severo Ochoa accreditation CEX2021-001142-S/MICIU/AEI/10.13039/501100011033. The funders had no role in study design, data collection and analysis, decision to publish, or preparation of the manuscript.

